# Does human primary motor cortex represent sequences of finger movements?

**DOI:** 10.1101/157438

**Authors:** Atsushi Yokoi, Spencer A. Arbuckle, Jörn Diedrichsen

**Affiliations:** The Brain and Mind Institute, University of Western Ontario, Canada; Graduate School of Frontier Biosciences, Osaka University, Japan

## Abstract

Human primary motor cortex (M1) is an essential structure for the production of dexterous hand movements. While distinct sub-populations of neurons are activated during single finger movements, it remains unknown whether M1 also represents sequences of multiple finger movements. Using novel multivariate fMRI analysis techniques, we show here that even after 5 days of intense practice there was little or no evidence for a true sequence representation in M1. Rather, the activity patterns for sequences in M1 could be explained by linear combination of patterns associated with the constituent individual finger movements, with the strongest weight on the finger making the first response of the sequence. These results suggest that M1 only represents single finger movements, but receives increased input at the start of a sequence. In contrast, the reliable differences between different sequences in premotor and parietal areas could not be explained by a strong weighting of the first finger, supporting the view that these regions exhibit a true representation of sequences.

## Introduction

Primary motor cortex (M1) with its direct projection to spinal motoneurons is a critical structure for fine hand control (Lawrence and Kuypers, 1968; Muir and Lemon, 1983). Population of neurons in M1 involved in individuated finger movement show considerable overlap (Schieber and Hibbard, 1993). Yet, they form large enough clusters to be detected with functional magnetic resonance imaging (fMRI) as unique activation pattern associated with each individual finger (Indovina and Sanes, 2001; Ejaz et al., 2015). Each of these populations can be conceptualized as a dynamical system (Churchland et al., 2012) (illustrated by arrows inside the two circles in Fig. 1A), that produces the continuous sequence of muscle activities necessary for the movement of a single finger. Here we ask whether such sub-populations in M1 can also learn to represent longer sequences that span movements of multiple different fingers.

**Figure 1.**
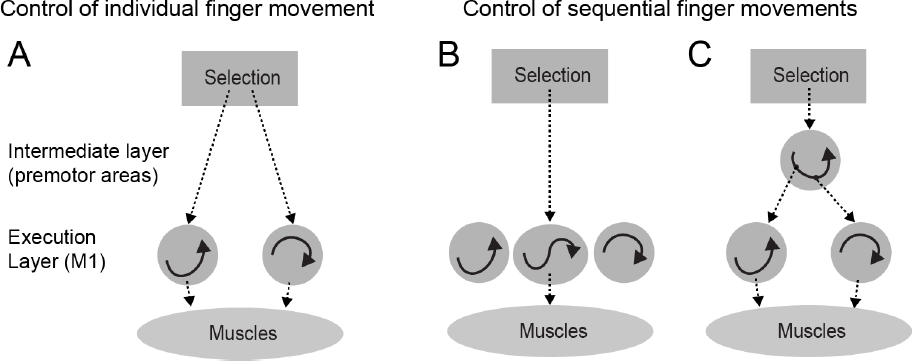
Two ways of representing sequential movements. (A)Before training sequences are produced through sequential selection of single finger movements. The execution layer (M1 and spinal cord) contains populations of neurons that, once activated, generate the muscle activity patterns necessary for a single finger movement through their intrinsic population dynamics. (B) After learning, the repeated sequential activation of two execution primitives leads to the formation of a new population of neurons that produces the two presses as a single unit. (C) Alternatively, a neural population in premotor areas could activate the execution primitives for the two fingers in the correct order.

Recent computational work has demonstrated that a randomly connected recurrent neural network can learn and store multiple dynamically evolving patterns (Laje and Buonomano, 2013). It is therefore conceivable that M1 develops dedicated populations of neurons that encode the sequences of two or more finger movements (Fig. 1B). In this scenario, the neural activity for pressing the 2^nd^ digit would be different depending on whether it was executed in the sequence 1-2 or 3-2. Such a representation would be necessary if M1 was to autonomously generate the spatio-temporal activity pattern necessary for sequence production. Indeed, it has been suggested that M1 acquires such representations of finger movement sequences after multiple days of training (Karni et al., 1995).

Alternatively, the learned sequences could be represented in secondary motor areas (Hikosaka et al., 2002; Diedrichsen and Kornysheva, 2015), which then activate the corresponding execution-related populations in M1 (Fig. 1C). A number of recording studies have found evidence of neurons that are uniquely activated for different sequences in dorsal premotor cortex (PMd) and the supplementary motor area (SMA) (Mushiake et al., 1991; Shima and Tanji, 1998). In this scenario, M1 would have no true sequence representation, as the neural activity would solely reflect the ongoing elementary movement independent of the sequential context (Mushiake et al., 1991; Ashe et al., 1993).

Here we sought to distinguish between these two possibilities, by analysing the fine-grained activity patterns in M1 using functional magnetic resonance imaging (fMRI) during the performance of well-learned finger sequences. Sequences consisted of different orderings of the same fingers presses. Because of the low temporal resolution of fMRI, we could not resolve the activity related to the individual presses, but could only measure the activity pattern averaged over the whole sequence. Nonetheless, if activity in M1 represented the movement sequence, we should find reliable differences between the sequences, as each sequence would activate a partly separate neuronal subpopulation (Fig. 1B).

Indeed, in previous studies (Wiestler and Diedrichsen, 2013; Kornysheva and Diedrichsen, 2014; Wiestler et al., 2014), we had found that sequences consisting of different permutations of the same five fingers can be reliably decoded from M1. However, the finding of decodeability alone does not provide unequivocal evidence for a true sequence representation. It is possible that activity in M1 only represents individual movements (i.e. that the activity for the second press is the same whether it is executed in the sequence 1-2 or 3-2, Fig. 1C), but that the amount of activity for each individual finger press depends on the serial position in the sequence. For example, it is possible that the first finger press in the sequence always elicits more activity than subsequent presses. Because we can only observe a temporally integrated signal in fMRI, such unequal weighting would lead to differences in activity patterns between different sequences. Thus to show evidence for a true sequence representation, we not only need to show distinguishable activity patterns for different sequences, but also demonstrate that these differences cannot be explained by a weighted combination of the activity patterns for individual presses.

To test this idea, we compared the patterns for multi-finger sequence with those obtained for the execution of repeated presses of each single finger. We found that in M1 differences between the activity patterns for different sequences could be fully explained by the combination of activity patterns elicited by single finger presses. Specifically, sequence activation patterns in M1 reflected a stronger activation for the first finger in the sequence than subsequent fingers. In contrast, activation patterns in premotor and parietal cortices could not be explained by a combination of the activity patterns for the elementary movements. This suggests that premotor areas comprise representations of movement sequence, which then activate the representations of the individual component movements in M1 (Fig. 1C).

## Results

### M1 “encodes” both single finger movements and sequences

We tested if sequences are represented within M1 by comparing the fine-grained fMRI brain activation pattern associated with fast finger sequences (6 finger presses within 2.5 sec) with those associated with single-finger movements. Participants practiced six sequences that comprised all orders of pressing the thumb, middle and little finger with their right hand (Fig. 2A,B). They also produced six repetitions of the same finger press with each of the fingers, as constituents of the sequences. Participants were trained for 3 days, approximately 6 hrs in total, until they could perform all sequences from memory without error and at the same speed. We localized areas that showed reliable differences between either single-finger or multi-finger sequences by using a surface-based search-light approach (Oosterhof et al., 2011). Based on previous results (Wiestler and Diedrichsen, 2013), we expected that different single finger movements and different movement sequences would elicit differentiable activity patterns in M1.

**Figure 2.**
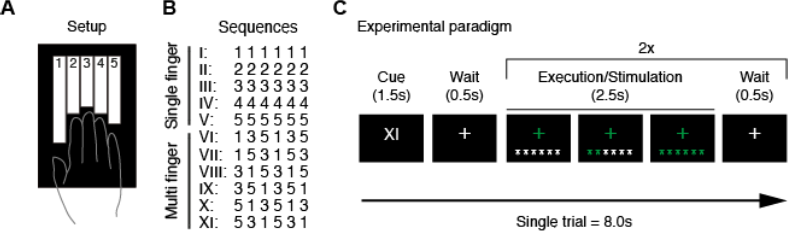
Methods for Experiment 1. (A) Participants generated isometric finger presses on a custom-built keyboard with force transducers and pneumatic pistons embedded within each key. (B) Participants were trained on five single-finger and 6 multi-finger sequences. (C) Schematic illustration for a trial during scanning. A roman numeral indicated the sequence to be executed. Participants then executed the sequence twice, receiving online visual feedback for each correct press. fMRI activity measurements were averaged across the two executions of the sequence, thereby removing temporal information from the activity profiles.

To characterize the representation, we calculated the cross-validated Mahalanobis distance (Walther et al., 2016) between the activity patterns for different conditions. As expected, we found evidence for a representation of single fingers in the hand area of primary motor (M1) and somatosensory cortex (S1, Fig 3A). Consistent with previous studies (Wiestler et al., 2011; Diedrichsen et al., 2013; Ejaz et al., 2015), weaker differences between activity patterns of single finger movements were also found in secondary motor areas such as dorsal and ventral premotor (PMd and PMv), supplementary motor (SMA) areas, and in the anterior superior parietal lobules, (aSPL, for stats see Figure 3-Supplement A,B), and the ipsilateral hemisphere (Diedrichsen et al., 2013).

**Figure 3.**
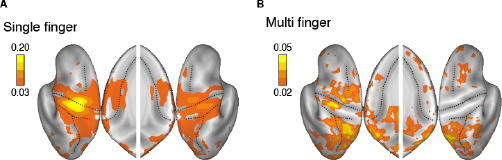
Searchlight map for movement representation. (A) Averaged distance for single finger sequences. (B) Averaged distance for multi finger sequences. Results are shown on an inflated view of the left and right hemisphere, with the inset showing distance on the medial wall.

The multi-finger sequences elicited differentiable activity patterns in premotor and parietal areas (Wiestler and Diedrichsen, 2013; Kornysheva and Diedrichsen, 2014; Wiestler et al., 2014) (Fig 3B). Importantly, we also found significantly different activity patterns for different sequences in M1 and S1 (Figure 3-Supplement C). The pattern distances for sequences were only 19±9% of those for single-finger movements, but they were reliable enough to decode which of the six sequences was performed with a cross-validated accuracy of 25±5% (chance-level is 16.67%).

One may argue that, if M1 only represented the individual finger presses, the activity patterns for the different sequences should have been indistinguishable. However, this argument relies on the assumption that all component actions elicit the same amount of activation regardless of the order in which they were made. Before concluding that M1 exhibits a genuine sequence representation (i.e. is in a different neuronal state for each sequence), we therefore need to consider the possibility that the input from premotor areas (Fig. 1C) varied depending on whether the finger press was in the beginning or middle of the sequence. As different sequences start with different fingers, this effect could lead to distinguishable BOLD activity patterns for different sequences, without implying a true representation of the sequence in M1.

### Differences in sequences depend on the first finger

To test for this possibility, we systematically compare the activation patterns for the multi-finger sequences to those of the single-finger presses and in M1. We calculated the cross-validated distances between all pairs of conditions in an anatomically defined region-of-interest (ROI; Figure 3-Supplement A) for contralateral M1. The resultant matrix of pair-wise distances (55 pairs in total) – the representational dissimilarity matrix (RDM) effectively summarizes the representational structure of the whole ROI (Fig. 4A).

**Figure 4.**
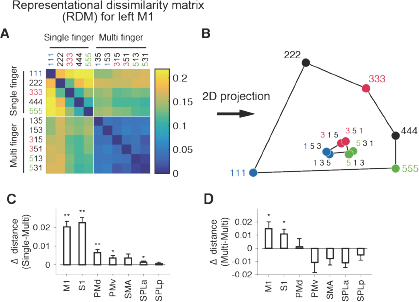
Representational structure in the left M1. (A) Representational dissimilarity matrix (RDM) calculated from the activation pattern within left M1 (contralateral to the performing hand). (B) Low-dimensional projection of the RDM by multi-dimensional scaling (MDS). Each dot represents a movement condition (1-5: single finger, 135-531: multi finger). (C, D) Test for first-finger effect (see Materials and Methods). (C) Mean distance between the single-finger movement (1, 3, or 5) and the multi-finger sequence that starts with the same finger MINUS the distance between the same single finger movement and sequences that start with a different finger. A positive difference indicates that the pattern for each multi-finger sequence is weighted towards the pattern of the first finger. (D) Mean distance (calculated for M1) between two multi-finger sequences that start with different fingers MINUS the mean distance between fingers that start with the same finger. A positive difference indicates that difference between sequences can partly be explained by the difference between the first finger. Asterisks indicate statistical significance assessed by onesided paired t-test (*: p<0.05, **: p<0.01).

To obtain insight into the representational structure, we applied a dimensionality reduction to the RDM by projecting it into a 2-dimensional space (Fig. 4B, for detail, see Materials and Methods). For single-finger movements (111, 333, etc.) we replicated the characteristic representational structure with the thumb showing the most unique pattern and the other fingers arranged in a semi-circle (Ejaz et al., 2015).

The multi-finger sequences are arranged such that two sequences starting with the same finger are clustered together (shown in the same colour in Fig. 4B). Furthermore, among all multi-finger sequence patterns, each pattern was also the most similar to the pattern associated with the first finger in the sequence. It should be noted, however, that low-dimensional projections (here designed to maximize the distances between single-finger movements, see Materials and Methods) do in general not capture all the aspects of the representational structure. Therefore, to fairly quantify these two key observations, we compared the cross-validated distances between activity patterns of single-and multi-finger sequences in the (un-projected) high-dimensional space.

If the activity pattern of each sequence is the most similar to the starting finger, then the pattern for each individual finger should be closer to sequences starting with this finger than to other sequences. For instance, the pattern for the thumb (111, in Fig. 4B) should be closer to sequence 135 and 153 than to other sequences (e.g., 351 or 315). This was indeed the case in contralateral M1 (*t*_8_=6.18, *p*=1.3×10^−4^) and S1 (*t*_8_=4.09, *p*=0.0018) (Fig. 4C).

Furthermore, two sequences starting with the same finger should be more similar to each other than other pairs (e.g., the distance 135 vs. 153 should be smaller than 135 vs. 315). Again, this effect was significant in M1 (Fig. 4D, *t*_8_=2.87, *p*=0.0104) and S1 (*t*_8_=3.08, p=0.0075). In contrast, no other tested ROI showed significance on both tests simultaneously (Fig. 4D).

One possible scenario which can explain both observations is that the activity patterns for sequences in M1 are a weighted sum of patterns elicited by the constituent single-finger presses, with the first finger having the highest weight. This would imply that there is no true sequence representation in M1. To evaluate whether this simple idea could fully explain the pattern differences between the multi-finger sequences in M1, we tested different candidate models for the activity patterns elicited by sequences using the framework of pattern component modeling (PCM) (Diedrichsen et al., 2011; Diedrichsen and Kriegeskorte, 2016; Diedrichsen et al., 2017). PCM allows us to directly compare different models for the representational structure inherent in the pattern of multi-voxel activities. Importantly, we can compare the model likelihood to a noise ceiling, to assess whether the model can fully account for the data given the level of measurement noise and inter-subject variability (see Materials & Method).

As a starting point, we assessed the “first-finger” model, in which the patterns for multi-finger sequences are the weighted sum of the single finger presses, with the first finger having the highest weight and all the subsequent fingers a lower, but equal, weights (see Materials & Method). This model predicts that sequences that start with the same finger do not have different patterns (Fig. 5A). We found that this model could almost fully account for the representational structure found in M1: the difference in log-likelihood to the Null-model (log-Bayes factor, see Materials & Method) fell between the upper and lower bound of the noise ceiling (Fig. 5B: 6.95 vs 6.45).

**Figure 5.**
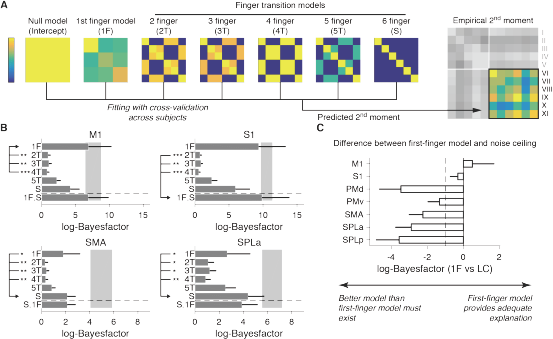
Evaluating representational models of multi finger sequences. (A) The empirical second moment of the activity patterns is modelled using a combination of the predicted second moment matrices for each of the models. (B) Difference in log-likelihood as compared to the null-model (log-Bayes-factor) for each component model. The grey area demarks the upper and lower noise ceiling. The combination of the first-finger model and the 6-finger transition model (1F.S) is shown below the horizontal dashed line. Winning model is marked by the arrow. Significant differences (assessed by Wilcoxon’s rank sum test on individual log-Bayes-factors) in the fit between the winning and the other models are marked by asterisks (*: p<0.05, **: p<0.01, ***: p<0.001). Error bars represent SE across subjects. (C) Log-Bayes-factor of the first-finger model compared to the lower noise ceiling for each ROI. Dashed line shows the typical threshold value for model selection (e.g., Kaas and Raftery 1995).

We then tested whether M1 might represent movement transitions between 2 or more fingers (see Fig 5A). Note that a representation of a 6-finger transition would mean that each sequence would have a unique activity pattern. The log-Bayes factor for these models was clearly lower than that for the first-finger model (Fig. 5B), indicating a poorer fit of these models.

We then explored linear combinations of models. Because the relative weight of each component was an additional free parameter, we evaluated the model likelihood using cross-validation across participants (see Materials & Methods). When we combined the first-finger model with the sequence model, we achieved a slightly lower likelihood than the first-finger model alone for M1, the average log-Bayes factor reduced by 0.05. For S1, however, the addition of sequence model achieved a slightly higher likelihood (9.37 for first-finger model alone, vs 9.87 for combined model, Fig. 5B). However, on a common scale of Bayes factors (Kass and Raftery, 1995), such a small difference would be considered *“not worth more than a bare mention”.*

In premotor areas, on the other hand, the representational structure was not well explained by the first-finger model. For example, in SMA and SPLa, the fit of the sequence model was systematically better than the first-finger model (Fig. 5B, for other ROIs, see Figure 5-supplement), indicating that the activity patterns in these regions represented sequential information. Importantly, the likelihood of the first-finger model was systematically below the lower bound of the noise ceiling (Fig. 5C): The mean difference in log *BF* to the lower noise ceiling was substantially larger than 1, indicating strong evidence (Kaas and Raftery, 1995) that for these regions a better model exists.

In summary, on the group level our results provided very limited evidence for a true, unique sequence representation, or the representation of transitions between fingers in M1. Instead, the representational structure for sequences in this area could almost fully be explained by the first-finger model – i.e assuming that the patterns for multi-finger sequences are a linear combination of the patterns associated with the individual finger presses, with the first finger weighted more strongly than the others. The same observation held true for S1. In contrast, in premotor regions the first-finger model could not account for the differences between sequences, suggesting genuine encoding of sequential information in these regions.

### First-finger effect in M1 is related to neural planning and execution processes

We hypothesize that that the prominent activity for the first finger press in M1 is related to active planning and execution processes. Given that the BOLD signal more closely reflects synaptic input than spiking activity of output neurons (Logothetis et al., 2001), one possible explanation is that M1 receives strong input from premotor regions at the beginning of the sequence to push the neural state from resting to active state at movement initiation. While M1 would still rely on premotor input to produce the subsequent finger presses, the amount of this input would be smaller as M1 is already in an active state.

Alternatively, the prominence of the first finger pattern could be due to the passive properties of M1. Specifically, the effect could have hemodynamic rather than neuronal causes. That is, the neural activity for each finger in the sequence could be exactly the same, but because of the non-linear integration of the BOLD signal for inter-stimulus intervals of <6s (Dale and Buckner, 1997), it may be that the first finger press achieved the majority of the vasodilatory response and hence dominates the overall activity pattern.

To rule out this possibility, we exploited the fact that the single-finger patterns in M1 and S1 can also be elicited by passive stimulation (Wiestler et al., 2011). In the scanner, we therefore “replayed” the recorded force traces during the active trials through pneumatic pistons mounted under each finger. If we can elicit comparable single-finger activity patterns in M1 through both active and passive movements, and if the timing of the presses is identical across conditions, then any hemodynamic, or passive neural effect, should apply equally in both situations. Thus, if the first-finger effect is due to the non-linear translation from neural to BOLD signals, we should find a similar representational structure for active and passive multi-finger movements.

As can be seen from Figure 6A, the spatial distribution of single finger representations was comparable to that obtained in the active condition (Fig. 3A). For a direct comparison, we calculated the average distances in each of the cortical ROIS (Fig. 6C). The distance in M1 was 82±11% of what was elicited in active condition, and 101±10% in S1. Additionally, the elicited patterns matched the active patterns on a finger-by-finger basis. The average correlation between active and passive patterns (after subtracting out the mean activity pattern) of the same finger were r=0.76±0.37, p=8.86 ×10^−5^, and r=0.89±0.05, p=6.8×10^−11^, respectively for M1 and S1. Therefore, we confirmed that almost comparable single-finger activity patterns are elicited in M1 through the passive stimulation.

**Figure 6.**
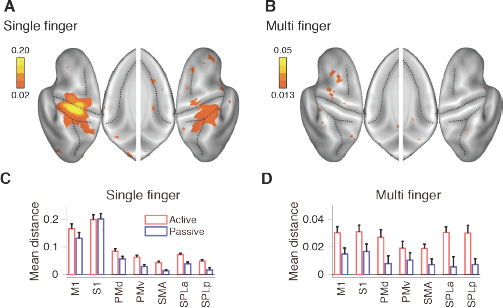
Passive stimulation elicited comparable single finger representation, but reduced multi-finger sequence representation. (A) Averaged distance for single finger sequences, and (B) multi finger sequences shown on an inflated view of the left and right hemisphere, with the inset showing distance on the medial wall. Scaling for the multi finger sequences were determined based on the reduction of distance from active movement condition for single finger sequences. (C) Average distance in the cortical ROIs (see Fig. 3 supplement) for single finger sequences for active (red) and passive (blue) conditions. (D) Average distance across all pairs for multi-finger sequences. Error bars represent SE across subjects.

In contrast to single-finger representations, encoding of multi-finger sequences reduced dramatically over the whole cortical surface (Fig. 6B). The distances between sequences reduced to 47±29% in M1 and 42±42% in S1 compared to the active condition (Fig. 6D). Critically, the reduction was larger than what would be expected from the reduction in the single-finger representations (Fig. 6B, M1: *t*_16_=1.7601, *p*=0.049, and S1: *t*_16_=2.587, *p*=0.001). If the first-finger effect had been solely due to a hemodynamic non-linearity, or to a passive adaptation of neural activity, then the effect should have equally applied to both active and passive conditions. Instead, the differences between active and passive conditions indicate that the high weighting of the first-finger press in M1 is caused by active preparation or initiation of the sequence.

The results also show that the sequence representations found in premotor regions are due to the active planning and execution of a sequence, and not to processing of the sensory inflow. The distances for multi-finger movements were substantially lower (24% on average) in premotor regions (Fig 6B) and not significantly different from zero in 4 of the 5 premotor ROIs. Furthermore, the remaining representational structure was relatively inconsistent between subjects, as can be seen in the low noise ceiling of the model fits (Supplemental Fig 6). These findings clearly indicate that the sequence representation observed in premotor regions requires the active execution of a sequence.

### A sequence representation with longer training?

So far, we have found little or no evidence for a real sequence representation in M1. We considered two reasons for this failure. First, it may be that the training period was too short. Secondly, the simple structure of our sequences (i.e. permutations of digit 1, 3, and 5) may have reduced the chances of forming representations of finger transitions.

We therefore conducted a second experiment, this time with 5-6 days of training of 2 hrs each, during which participants learned 8 arbitrary sequences, each 11 presses long consisting of all five fingers. The trained sequences were executed at preferred speed (average 4.3 presses / s) in the scanner (see Material & Methods).

First, due to the fact that experiment 2 had more data, the evidence for movement representation was substantially stronger. The scale of log-Bayes factor was approximately 3~4 times larger in Figure 7 compared with Figure 5. However, despite increased signal and despite ample opportunity to form representations of at least two or three finger transitions, the representational structure in M1 was again fully explained by the first finger model. The log-Bayes factor of the first-finger model was above the noise ceiling. Addition of the sequence representation or a 2-finger transition model did not substantially improve the fit (Fig. 7A, 18.95 vs 19.16 for single model and combined model, respectively). Similar result was obtained for S1. However, in this region the addition of the sequence model slightly improved the likelihood of the model (Fig. 7A, 45.84 vs 47.84 for single model and combined model, respectively). In contrast, the representational structure in premotor and parietal regions could not be explained by the first-finger model, suggesting the presence of a more complex and higher-level sequence representation (Fig. 7B). The content of these representations, and their dependence on cognitive mechanisms of movement chunking, will be reported in a subsequent paper. For M1, however, these results confirm that even after week-long training, the activity pattern reflect processes related to the individual finger presses, but not to their sequential context.

**Figure 7.**
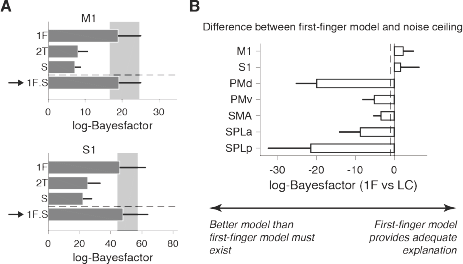
More intensive training with complex sequences (Experiment 2) revealed highly similar results. Participants in the second experiment practiced 8 different sequences of 11 presses-long for 5 days and 2 hrs per a day before the imaging session. (A) log-Bayes-factor (Fig. 5) for the data of M1 and S1 for the first finger model (1F), the 2-finger transition model (2T), sequence model (S) and the combination between first finger and sequence model (1F+S). The arrow again indicates the winning model. Error bars represent SE across subjects. (B) Log-Bayes factor between first-finger and noise ceiling model. As in Experiment 1, the first-finger model provides an adequate explanation for M1 and S1, but not for secondary motor areas. Dashed line shows the typical threshold value for model selection (e.g., Kaas and Raftery 1995).

## Discussion

We demonstrated that even after 5-6 days of intensive practice, there was very little evidence for a genuine sequence representation in M1. We also did not find evidence for a representation of partial sequences, such as the transition between 2 or more finger presses. Instead, we found that the activity patterns for sequences could be explained by a linear combination of the activity patterns for single finger presses, in which the weight of the first finger was higher than for the other presses. This resulted in an above-chance classification accuracy for sequences beginning with different fingers. We also provided evidence that this first-finger effect was much larger during active compared to passive sequence production, arguing that it is related to active movement preparation and initiation. These results also indicate that the first-finger effect had a neural origin, rather than being based on a hemodynamic non-linearity. In contrast to the absence of any sequential information in M1, sequences were robustly represented in secondary motor areas, such as PMd, SMA, and the anterior SPL. These areas have been shown to represent sequences such that the neural activity pattern reflects the sequential order of movement elements (Mushiake et al., 1991; Tanji and Shima, 1994; Wiestler and Diedrichsen, 2013; Wiestler et al., 2014).

### Advances from the earlier studies

Although there have been numerous imaging studies on sequence production and acquisition (Karni et al., 1995; Honda et al., 1998; Karni et al., 1998; Doyon et al., 2002), our approach makes several advances over these earlier studies. First, we designed our experiment specifically for multi-voxel pattern analysis, which allowed us to test directly for sequence representations in M1. This is not possible when looking only at the average BOLD activity within a region. Indeed, the increases and decreases reported in previous studies (Grafton et al., 1995; Karni et al., 1995; Honda et al., 1998; Karni et al., 1998; Kawashima et al., 1998; Sakai et al., 1998) may not necessarily reflect plastic changes in M1. Rather, they could equally well reflect changes in the input to M1, caused by sequences being learned and represented in secondary motor regions. Multi-voxel pattern analysis is also sensitive to inputs from other regions, but reveals the local organization of how these inputs arrive in M1. Specifically, our results suggest that the activity patterns for the first finger press are especially high, but that the underlying activity patterns still only reflect the individual finger movements.

Second, we not only measured the activation pattern for the sequences, but also compared them to the patterns of their constituent single finger movements. This allowed us to determine whether the activity patterns for multi-finger sequences could be explained by a combination of single-finger movements, or whether there was evidence for a new representation that encoded the sequential context (Fig 1B). Our results clearly argue for the former, implying that the significant differences between sequence patterns in M1 in our earlier work (Wiestler and Diedrichsen, 2013; Kornysheva and Diedrichsen, 2014; Wiestler et al., 2014) did not reflect an encoding of the order of finger presses (i.e. a genuine sequence representation), but of the sequential position of finger presses. In these studies, the different sequences started with a different finger, such that we could not distinguish a real sequence representation from one caused by the first-finger effect. Importantly, our current result confirmed that the pattern differences reported in secondary motor areas reflect genuine sequence encoding.

Finally, we demonstrated that in secondary motor areas no robust sequence representation could be elicited using passive sensory stimulation. This suggests that the sequence representations observed in these areas actually reflects active movement planning/execution process, rather than sensory re-afferent signals. Of course, the sensory feedback during the passive stimulation condition was not exactly the same as during active sequence production. However, nearly identical activity patterns in the single-finger conditions elicited in primary sensory cortex by the passive stimuli demonstrated that the sensory feedback closely mimicked that during active presses.

### The origin of the first-finger effect

The results from the passive stimulation also argue that the first-finger effect is related to the active preparation and initiation of the sequence, rather than just to the sensory inflow. More generally, the results show that the effect has a neural origin, and is not purely caused by a non-linear integration of neural events in the production of the hemodynamic response (Dale and Buckner, 1997). Recent electrophysiological findings seem to support this conclusion (Hermes et al., 2012; Siero et al., 2013). These studies recorded the electrophysiological potentials using intracranial ECoG electrodes above M1 while human participants performed rhythmic open and close movements of hand at ~2Hz. Power in the high-gamma frequency band was much more pronounced for the first movement of a sequence as compared to subsequent movements (Hermes et al., 2012). Siero et al. (2013) also showed that the high-gamma activity tightly related to the observed BOLD activity recorded when subjects performed the same task in the scanner. While the “sequence” in these experiments consisted of the repetition of the same movement elements, our results lead to the prediction that similar effect should occur for more complex, multi-finger sequences.

What is the neural origin of this first-finger effect? First note that both BOLD and high-frequency gamma power relate mainly to synaptic input to a region. Thus, it is not unlikely that this effect arises only on the input side and that the firing of output neurons would be matched for the different finger presses (Picard et al., 2013). The most likely explanation therefore is that the neural circuits in M1 require a large input drive to initiate a series of movements. Recent results have shown that the largest change in neural activity occurs when transitioning between a “resting” sub-space to the active sub-space (Elsayed et al., 2016). In our case the driving input for this movement would arrive in form of the intention to move the first finger. Subsequent finger presses would still require input from higher-order areas, as M1 would not be able to generate the sequence autonomously, but the input drive would be much smaller as the state of the neurons would already be in the vicinity of the active subspace. This idea also predicts that if the sequence is executed slowly enough, the state in M1 should relax back to the resting sub-space and the first-finger effect should disappear.

### Limitations: length of training and sequence representation in M1

Our data provides very little or no evidence for a sequence representation in M1 after 1 week of intensive training (1.5-2 hours per a day). However, this does not exclude the possibility that longer period of training might result in the unique neural circuits for sequences acquired within M1. After 2 years of training, a single-cell recording study in the monkey revealed some evidence for sequential representations in M1 (Matsuzaka et al., 2007). Note however, that in this study, sequence representations were assessed as the difference between neuronal responses to trained and untrained sequences, not as in our study between different trained sequences. On a much shorter time scale, Karni et al. (1995) reported an expansion of activated area in M1 over 4 weeks of daily practice. The total amount of practice was similar for the experiments reported here (approx. 3.5~7 hrs vs. 6~10 hrs in our study). Again, the results only indicated that trained sequences elicited more activity than untrained sequences (a result that we failed to replicate, Wiestler and Diedrichsen, 2013), but does not show the presence of neural processes that would relate to the sequential order of movement elements.

Using our methods, we did not find evidence for the representation of short sequence components, such as the transition between 2 or 3 fingers. There was some indication that there was a weak component of the activity pattern in M1 which may reflect the sequence itself. We are now investigating whether these patterns constitute the beginning of a “true” sequence representations that will increase in strength with extended training.

## Conclusion

Using representational fMRI analysis, we demonstrated that up to about 1 week of intensive practice, activity in M1 relates to individual finger presses, but not to transitions between multiple fingers or even full sequences. At the same time, we found robust and genuine sequence representation in other higher motor areas, such as PMd, SMA, or aSPL, which is consistent with previous studies (Mushiake et al., 1991; Shima and Tanji, 1998). The next challenge is to dissect the content of these representations in detail (Lashley, 1951).

## Materials & Methods

### Participants

Nine healthy, right-handed volunteers (3 females, age: 23±4) participated in Experiment 1, and 14 healthy, right-handed volunteers (8 females, age: 23±3) participated in Experiment 2, after providing written informed consent. The experimental procedures were approved by local ethics committees at the University of Western Ontario (London, Canada) and University College London (London, UK). None of the participants was professional musician nor has any known neurological history.

### Apparatus

We used custom-build five-finger keyboards (Fig. 2A) with a force transducer (Honeywell FS series) mounted underneath each key (Wiestler and Diedrichsen; Wiestler et al.). The keys were immobile and measured isometric finger force production. Dynamic range of the force transducers was 0-16N and the resolution <0.02 (N). A finger press/release was detected when the force value crossed a threshold of approximately 3 N. This threshold was slightly adjusted for each finger to ensure that each key could be pressed easily. The signal from the keyboard were low-pass filtered, amplified and sent to PC for online task control and data recording. The forces were recorded at 200 Hz. For passive stimulation of the fingers, a pneumatic air piston was mounted underneath each key. The pistons were driven by compressed air (100 psi) from outside the MRI scanning room through poly-vinyl tubes. The force exerted by each piston was controlled by a pressure-regulating valves. The movements of the fingers was restricted by a device mounted above the fingers.

### Sequence production task for Experiment 1

During the training sessions, participants were seated in front of the LCD monitor and placed their fingers on the keyboard. They learned to produce five single finger sequences and six multi-finger sequences. For the single-finger sequences, one of five fingers had to be pressed 6 times (e.g., 3 3 3 3 3 3); for the multi-finger sequences one of the six possible permutations of fingers 1, 3, and 5 was pressed twice (e.g., 5 3 1 5 3 1) (Fig. 2B). All fingers remained on the keyboard at all times, such that the overt movement of the fingers was minimized.

The participants practiced the sequences for 3 days so they were able to produce the sequences in the scanner within 2.5 seconds from memory given only visual cue, which was presented for 1.5 seconds at the start of each trial (Fig. 2C). Each sequence was indicated by a different Roman numeral (I, II, …, XI). In the beginning of training we provided both the sequence cue (roman numeral) and all six to-be-pressed digits on the screen. Subsequently, we replaced the digits with asterisks (*), to encourage the participants to memorise the sequences (Fig. 2C).

A total 1716 sequence executions were made (156 executions per one sequence type). The order of 11 sequences was pseudo-randomised throughout the sessions. The colour of a asterisks turned to green immediately after a press was correctly registered, while it turned to red if the press was incorrect. To guide participants’ speed, the sequence cue blinked at a reference frequency that gradually increased during the training sessions at constant rate until it reached to 4 Hz. On the last day of training sessions, participants practiced actual task for the scanning session, lying on the mock MRI scanner bed for familiarisation.

### Sequence production task for Experiment 2

The general methods were similar to the first experiment. Participants learned to produce 8 different sequences with 11 presses from the memory. Initially we trained participants for 5 days, but for the other half added a 6^th^ day, such that all could correctly produce the sequences within 2.5 seconds. On average, the training lasted cumulatively 10-12 hrs. As in Experiment 1, the sequences were cued with Roman numerals I–VIII. All the sequences were matched with the number of finger presses used; 2 presses with thumb, middle, ring, and little fingers, and 3 presses with index finger, respectively. Four of the sequences started with the thumb, two sequences started with middle finger, and the rest of two sequences started with little finger. The detailed training protocol and the behavioural results of training and transfer test (conducted after the imaging) will be reported in a separate paper.

### Imaging session

During the imaging session, the participants lay supine on the scanner bed with knees slightly bent supported by a wedge-shaped cushion. The pneumatic keyboard was comfortably placed on their lap, and visual stimuli were presented on a back-projection screen which was viewed through a mirror attached to the head coil.

For Experiment 1, we conducted both active and passive conditions. In each trial of the active condition, the participants were first provided with the sequence cue for 1.5 seconds and then they were required to execute the specified sequence twice within the time limit of 2.5 seconds for each execution (Fig. 2C). Each execution was triggered by the fixation cross turning green. During the execution period, the fixation cross blinked at the reference frequency (4 Hz) to provide the participants with a pacing signal. The order of the 11 sequences was pseudo-randomised and included 1 rest trial of 8 seconds, during which the participants only passively viewed the fixation cross. This set of sequences was repeated three times within each imaging run, resulting in a total of 66 sequence executions per run. We conducted seven runs in the active condition. For these runs, there was also no significant difference in the pressing frequency (Hz) between single and multi-finger sequences (4.58±0.36, 4.59±0.39, *t*_8_=−0.176, *p*=0.865). The average number of incorrect presses per each execution was close to zero, but slightly larger for multi finger sequences (0.02±0.02, 0.22±0.13, *t*_8_=−4.884, *p*=0.001).

Alternating with the active runs, we conducted seven imaging runs in the passive condition. During the active run, we recorded the force data to replay these forces through the pistons in the passive run. The visual stimuli and timing were exactly the same as in the active runs, except that the participants were told not to produce any active finger movement, but to only passively receive stimulations to their fingers. Each passive run used the exact timings of the preceding active run, only that the sequence of trials was randomly shuffled on each run. Due to the nonlinear response property of pneumatic pistons, the resultant passive forces were lower than the forces in the active condition. We confirmed, however, that we could elicited robust single finger representation almost comparable to the active condition, especially in S1 (see Results).

The structure of Experiment 2 was similar. In the beginning of each trial, the sequence cue (I-VIII) was presented for 2.5 seconds. This was followed by two execution phases of 4 seconds each, with 0.5-second ITI. During the execution phase, only fixation cross and asterisks were presented. The order of sequences was randomised, and each of the 8 sequences was repeated three times during each run. During scanning the average pressing frequency was 4.47±1.05 Hz, which was not significantly different from that in the Exp 1 (*t*_19_ = 0.298, *p* = 0.769). Four resting trials of 10.5 seconds were randomly interspersed. We conducted a total of 9 runs, each of which lasted a total of about 7 min. Short breaks (up to a few minutes) were interleaved when subjects required. There was no passive condition for this experiment. The average number of incorrect presses per each execution was again close to zero, but significantly larger than that in the Exp 1 (0.40±0.16, *t*_19_<4.84, *p*=1.13×10^−4^).

### Imaging data acquisition

Experiment 1 was conducted on a Siemens Magnetom Syngo 7T MRI scanner system with a 32-channel head coil at the Centre for Functional and Metabolic Mapping, Robarts Research Institute (London, Ontario, Canada). Inhomogeneity of main magnetic field was adjusted by B0 and B1 shimming at the beginning of the whole session. Functional images were acquired for 14 imaging runs of 300 volumes per each using multi-band 2-D echo-planer imaging sequence (TR = 1.00 sec, multi-band acceleration factor = 2, in-plane acceleration factor = 3, resolution: 2.0 mm isotropic with 0.2 mm gap between slices, and 44 slices interleaved). The first 4 volumes were discarded to ensure stable magnetization. The slices were acquired close to axial to cover the dorsal aspects of the brain, including most of the frontal, parietal, occipital lobes, and basal ganglia. The ventral aspects of the frontal and temporal lobes, brainstem, and the cerebellum were not covered. Each functional imaging run lasted for 5 minutes. T1 weighted anatomical image was obtained on a separate session using MP2RAGE sequence (TR = 6.0 sec, resolution: 0.75 mm isotropic).

Experiment 2 was conducted on a Siemens Trio 3T scanner system with a 32-channel head coil at the Welcome Trust Centre for Neuroimaging (London, United Kingdom). B0-field maps were acquired at the beginning of the whole session to correct for inhomogeneity of main magnetic field (Hutton et al., 2002). Functional images were acquired for 9 runs of 135 volumes each using 2-D echo-planer imaging sequence (TR=2.72 sec, in-plane acceleration factor = 2, resolution = 2.3mm isotropic with 0.3 mm gap between each slice, and 32 slice interleaved). The first 5 volumes were discarded to ensure stable magnetization. The coverage was similar to Experiment 1. A T1 weighted anatomical image was obtained using MPRAGE sequence (1mm isotropic resolution).

### Behavioural data analysis

Recorded force data were analyzed offline. The data for both the training and scanning sessions was first smoothed with second-order Butterworth filter with cutoff frequency of 10 Hz to remove remaining RF noise and then submitted to the subsequent analysis. Press and release timings were defined as the time point where the press force first crossed the threshold (3 N) and then returned to the below-threshold level. Reaction time (RT) from the go cue, movement time (MT) starting from first press time to the last release time, inter-press intervals (IPIs), and the number of incorrect presses at each execution were calculated.

### Imaging data analysis

#### Preprocessing and first-level model

Experiment 1: Functional imaging data were pre-processed using SPM12 (http://www.fil.ion.ucl.ac.uk/spm/). Functional images were first motion corrected, and the mean images were co-registered to the individual anatomical image. As we had relatively fast TR (1.0 sec), we did not correct for slice acquisition timing. The data were then submitted to the 1st-level GLM to estimate the size of the evoked activity for each sequence type and run. We modelled the temporal autocorrelation using the “fast” option, which provides a flexible basis function to model dependencies on longer time scales. High-pass filtering was achieved by temporally pre-whitening the matrix using the temporal autocorrelation estimate.

*Experiment 2:* Pre-processing and GLM was conducted as in Experiment 1 – with the exception that we corrected for slice timing (given the slower TR). We also corrected for B0 inhomogeneity by using field map images. Given the slower TR, for the 1st-level GLM we used the standard high-pass filtering with a cut-off frequency of 128s and robust-weighted least square (RWLS, Diedrichsen and Shadmehr, 2005). The data from two participants in Experiment 2 was excluded from further analyses due to poor behavioural performance during scanning. These participants lacked a single correct trial in one of sequence types at more than one session. Hence, only the data from the remaining 12 participants were submitted to subsequent analyses.

#### Searchlight and ROI definition

Individual cortical surfaces (i.e., the pial and white-grey matter surfaces) were reconstructed from the anatomical image by using Freesurfer software (Fischl et al., 1999). We defined a continuous surface-based searchlight (Oosterhof et al.) as small circular cortical patches (approximately 11 mm radius) centred on each node defined on the reconstructed cortical surface that contains 160 voxels. Anatomical regions of interest (ROIs) were defined on this reconstructed surface (Fig. 3-SA) exactly as reported in previous studies (Wiestler and Diedrichsen; Kornysheva and Diedrichsen, 2014; Wiestler et al.).

#### Multivariate fMRI analysis

Within each of these groups of voxels (surface-based searchlight or anatomically-defined ROIs) we extracted the beta-weights for each sequence type and imaging run. We then spatially pre-whitened this the activity estimates across voxels in each area using multivariate noise-normalization with a regularized estimate of the overall noise-covariance matrix (Walther et al., 2016). This procedure renders the resultant voxels approximately uncorrelated in the noise (Diedrichsen and Kriegeskorte, 2016).

For these voxels, we then analyzed how the different multivariate activity patterns represented the sequences, using the representation model framework (Diedrichsen and Kriegeskorte, 2016). In this framework, the representational structure is described either by asking how the measured activity profiles of individual voxels are distributed in the space of experimental conditions, or – equivalently how distinguishable each pair of activity patterns associated with these conditions are from each other. Both viewpoints rely on a single central sufficient statistic, namely the second moment matrix of the activity patterns.

If **U** represents the true pattern of interest for the *K* experimental conditions times *P* voxels, then the second moment between the activity patterns is defined as

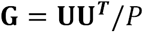

We analyzed this quantity using two complementary approaches: representational similarity analysis (RSA) to establish basic features of the representation and for visualization and Pattern component modelling (PCM) to compare more complex representational models.

#### Representational Similarity Analysis (RSA)

In RSA, we quantify the representational structure by measuring how distinct each pair of activity patterns are from each other. The squared Euclidean distance between the activity pattern **u**_1_ and **u**_2_ for example is:

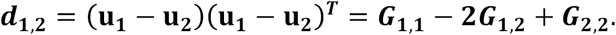

Calculated on spatially pre-whitened data, this distance is equal to the squared Mahalanobis distance. One problem is that estimates of this distance based on noisy data are positively biased. We therefore used here a cross-validated estimate of the second moment matrix **G**,

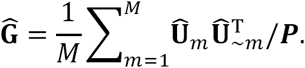

where *M* is the total number of partitions (e.g. imaging runs), **Û**_m_ is estimated prewhitened activity pattern for partition *m*, and **Û**_~*m*_ is the estimate of the pattern independent of the partition *m*. The “crossnobis estimator” (Diedrichsen and Kriegeskorte, 2016; Walther et al., 2016) is a distance calculated using this second moment matrix. This distance estimator is unbiased – meaning it can be used to directly test whether a distances is larger than zero. Finding consistently positive distance estimates therefor implies that the two condition activity patterns differ from each other more than expected by chance.

To visualize the representational structure, we used classical multi-dimensional scaling. We first projected the activity patterns into a lower dimensional sub-space by finding the Eigenvectors of group-averaged **Ĝ** matrix, which were then weighted by the square root of corresponding Eigenvalues. The projection displayed in Fig. 4B was then rotated to maximize the differences between the single-finger movements.

#### Pattern component modelling (PCM)

To compare full models of the representational structure, we used PCM (Diedrichsen et al., 2011; Diedrichsen et al., 2017), which directly evaluates the likelihood of the data under the linear model

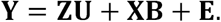

Here, **Y** is a *N*-by-*P* matrix representing noise-normalized activity pattern after the 1st-level GLM (Walther et al., 2016), where *N* is the number of estimates (number of conditions x number of runs) and *P* is the number of voxels. **Z** (*N*-by-*K* matrix) is the design matrix that associates **U** and **Y**. **B** represents the patterns of no interest, in our case the mean activity pattern in each run. Finally, **E** represents trial-by-trial measurement errors.

Importantly, PCM considers the true activity patterns **U** to be a random variable that follows multivariate normal distribution as, **U~N(0, G)**, where **G** is the second moment matrix of activity pattern **U**, which determines the similarity structure across movement conditions. In evaluating models, PCM integrates the actual activity patterns out, i.e. it evaluates the marginal likelihood (simply termed likelihood in this paper);

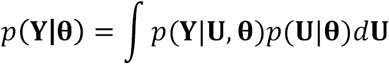

where **θ** represents model parameters that determine the shape of G and the signal and noise variances (see Diedrichsen et al., 2017). We fitted a number of models to explain the representational structure of the patterns associated with the multi-finger sequences.

*1st-finger model:* In this model, we assumed that the activity patterns for the multi-finger sequences are a weighted linear combination of the patterns for the constituent single finger presses. If all fingers were weighted equivalently, the overall patterns would identical, as each sequence contains exactly the same fingers. The 1st-finger model assumes that the first finger press is more strongly weighted than subsequent presses. Thus, the activity pattern for the multi-finger sequences are modelled as weighted sum of the activity pattern for the single-finger sequences,

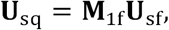

where **U**_sq_ is the pattern for multi-finger sequences (6×*P* matrix), **U**_sf_ is the activation patterns for the single finger presses of thumb, middle, and little fingers (3×*P* matrix), and **M**_1f_ is the weight matrix. Because each finger is present in each sequence equally often, we can simply model the difference in weight between the first and the subsequent fingers, such that **M**_1f_ is set to 1 for the first finger, and 0 otherwise (6×3 matrix). Therefore, the predicted similarity structure across multi-finger sequences (i.e., the second moment of the pattern **G**_i_f) is fully determined from the similarity across the single finger presses (i.e., **G**_sf_)

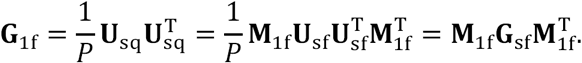

This results in the specific similarity structure depicted in Figure 5A. For modelling the activity at different ROIs, the empirical estimate of **G**_sf_ was derived for each ROI from the data – therefore no free parameter was required for this model.

*N-finger transition model:* This model family predicts the similarity structure based on neural circuits that encode the transitions between finger presses. Unique transitions can be defined between pairs of presses, or based on 3 or more presses. For instance, each sequence has five specific two-finger transitions, four three-finger transitions, etc. Thus, the predicted activity patterns of the multi-finger sequences are

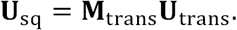

In this case the weighting matrix **M**_trans_ indicates, for each sequence, how many of the possible 2-digit transitions (9 total), 3-digit transitions (27 total), etc. the sequences contained, and **U**_trans_ represents specific activation patterns for each possible transition. Because we did not measure patterns for individual transitions, we assumed that each transition would be equally-strongly and independently represented, i.e., 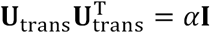, where *α* is constant and **I** is identity matrix. Thus, the predicted second moment matrix is

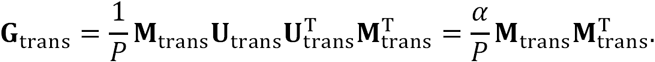

The resultant predicted similarity structure for each N-finger transition model can be seen in (Fig. 5A). Note that the six-finger transition model predicts that all sequences are equally distinct from each other, as each sequence has only one unique six-finger transition (the sequence).

#### Model comparison

We first fitted the six individual models (see previous section) separately. To account for individual differences in the signal-to-noise ratio, we maximized the likelihood in respect to a noise and signal strength parameter (Diedrichsen et al., 2017) – thus each model had the same two free parameters, allowing us to compare their likelihoods directly. We also fitted combinations of models, where the overall representation was a mixture of the hypothesized representations (i.e., the second moment matrix the weighted sum of those models). In this case, each component weight added an additional free parameter. Therefore, each single model has 2 free parameters (i.e., signal and noise parameters), and each mixture of two models has 3 free parameters (i.e., signal, noise, and the mixing ratio of one model over the other). Note that the *α* in the finger transition models is absorbed into the signal parameter.

To compare models with different number of parameters, we used group cross-validation: We fitted the parameters using the data from n-1 subjects, and then use the estimated G to fit the data from the left-out subject, fixing the parameters for G (for more details, see Diedrichsen et al., 2017). Note that in this process an overall signal and noise parameter was always fitted individually to each subject. Through this process, we obtained a cross-validated likelihood for each candidate models and subject, which serves as an estimate of the model evidence for each model.

We then compared models by calculating the log-Bayes factor which tells us to what degree one model can better describe the observed data over the other calculated (Hackett and Kaas, 2004) as the difference between the log-likelihoods:

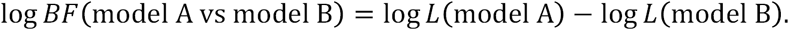

*logBF* were computed separately for each subject. We then used standard criteria for the average logBF proposed by Kaas and Raftery (1995) to judge if a model is meaningfully “better” than the other. Instead of using the group log-Bayes factor (Stephan et al., 2009), i.e. the sum of the individual log-Bayes factor, we report here the average logBF, which is invariant to the number of participants. This provides a much more stringent criterion for model selection.

### Noise ceiling

We also estimated the likelihood that the best achievable model should reach, called noise ceiling. The noise ceiling is an important measure to assess whether the selected model is a sufficient model, or whether the model misses a substantial aspect of the representational structure that is consistently observed across individuals. For this we used a free (fully flexible) model, which has as many parameters as the number of the elements in the second-moment matrix. For an estimate of the free model, we simply used the mean of cross-validated second moment matrix **Ĝ** across subject, which gives nearly identical results as using the maximum-likelihood estimate (for details see Diedrichsen et al., 2017).

To determine the free model, we first used the data of all subjects combined. This results in the best achievable likelihood for a group model and therefore constitutes an upper bound for the likelihood. Because this estimate is over-fitted, we also determined the cross-validated likelihood of the free model, which constitutes a lower bound estimate of noise ceiling. Therefore, even if a model performs better than the lower noise ceiling, it remains be possible that a better model still exists. However, based on the absolute performance we can conclude that the model captures all clearly consistent effects in the data.

## Statistics

We used one-sided, one-sample t-test for the evaluation of positive mean distance across subjects. To assess the first-finger effect, we performed two kinds of separate paired-t tests; a) if distances between two multi-finger sequences sharing the same first finger are smaller than distances between any other pair of multi-finger sequences not sharing the same first finger, b) if distances between a single-finger sequence and a multi-finger sequence sharing the same first finger are smaller than distances between any other pairs between single-finger and multi-finger sequences not sharing the same first finger. Significant difference for both of above comparisons (a and b) was deemed as the evidence of the first-finger effect. The ratio between active and passive distances (i.e., the reduction of passive distance) was estimated using linear-regression without intercept. Estimated slopes between single-and multi-finger sequences were then compared using simple t-contrast.

For the model comparison using PCM, we employed the standard interpretation of the size of the BF (Kaas and Raftery, 1995, see above. Additionally, we also report a Wilcoxon’s rank sum test on the log-Bayes factors between the winning and other models. Significance level was set to p=0.05. All the statistical analyses were performed on MATLAB (Mathworks, Inc.).

## Acknowledgements

Funding: JSPS Postdoctoral Fellowship (#15J03233) to AY, a James S. McDonnell Foundation Scholar award, and NSERC Discovery Grant (RGPIN-2016-04890) to JD. People: M Mohan for assistance in data collection, A Pruszynski and N Hagura for comments on early version of manuscript, and A Haith, M Smith, and R Ivry for comments and discussions on the manuscript.

## Open source programs

The MATLAB code used for the multivariate fMRI analysis (pattern component modelling) are available online (https://github.com/jdiedrichsen/pcm_toolbox).

---

**Figure 3-Supplement.**
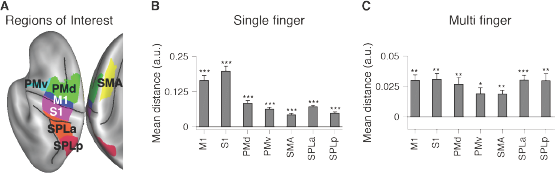
ROI analysis of movement representation. (A) Seven ROIs were defined on the left hemisphere of reconstructed cortical surface. (B,C) Mean distances calculated for single finger sequences (B), and multi finger sequences (C). Asterisks indicate significance based on the group t-test. *: p<0.05, **: p<0.01, ***: p<0.001.

**Figure 5-Supplement.**
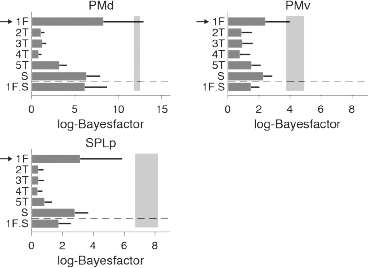
Model fitting result for other ROIs.

**Figure 6-Supplement.**
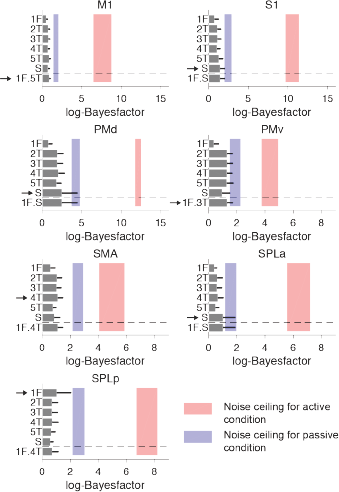
Model-fitting of multi-finger sequences for passive stimulation evaluated at each ROI. We applied the same model-fitting procedure as shown in Figure 5 to the data of passive stimulation condition. In contrast to the active movement case, the models performed almost equally poor for all ROI tested. Furthermore, group-wise consistency of representational structure (blue shaded areas, lower and upper noise-ceilings) was much lower compared with active movements (red shaded areas).

## References

Ashe J, Taira M, Smyrnis N, Pellizzer G, Georgakopoulos T, Lurito JT, Georgopoulos AP (1993) Motor cortical activity preceding a memorized movement trajectory with an orthogonal bend.

Churchland MM, Cunningham JP, Kaufman MT, Foster JD, Nuyujukian P, Ryu SI, Shenoy KV (2012) Neural population dynamics during reaching. Nature 487:51–56.

Dale AM, Buckner RL (1997) Selective averaging of rapidly presented individual trials using fMRI. Hum Brain Mapp 5:329–340.

Diedrichsen J, Shadmehr R (2005) Detecting and adjusting for artifacts in fMRI time series data. Neuroimage 27:624–634.

Diedrichsen J, Kornysheva K (2015) Motor skill learning between selection and execution. Trends in cognitive sciences 19:227–233.

Diedrichsen J, Kriegeskorte N (2016) Representational models: A common framework for understanding encoding, pattern-component, and representational-similarity analysis. BioRxiv.

Diedrichsen J, Wiestler T, Krakauer JW (2013) Two distinct ipsilateral cortical representations for individuated finger movements. Cereb Cortex 23:1362–1377.

Diedrichsen J, Yokoi A, Arbuckle S (2017) Pattern Component Modeling: A Flexible Approach For Understanding The Representational Structure Of Brain Activity Patterns. bioRxiv.

Diedrichsen J, Ridgway GR, Friston KJ, Wiestler T (2011) Comparing the similarity and spatial structure of neural representations: a pattern-component model. Neuroimage 55:1665–1678.

Doyon J, Song AW, Karni A, Lalonde F, Adams MM, Ungerleider LG (2002) Experience-dependent changes in cerebellar contributions to motor sequence learning. Proc Natl Acad Sci U S A 99:1017–1022.

Ejaz N, Hamada M, Diedrichsen J (2015) Hand use predicts the structure of representations in sensorimotor cortex. Nature neuroscience 18:1034–1040.

Elsayed GF, Lara AH, Kaufman MT, Churchland MM, Cunningham JP (2016) Reorganization between preparatory and movement population responses in motor cortex. Nature Communications 7:13239.

Fischl B, Sereno MI, Tootell RBH, Dale AM (1999) High-resolution intersubject averaging and a coordinate system for the cortical surface. Human brain mapping 8:272–284.

Grafton ST, Hazeltine E, Ivry R (1995) Functional mapping of sequence learning in normal humans. Journal of Cognitive Neuroscience 7:497–510.

Hackett TA, Kaas JH (2004) Auditory Cortex in Primates: Functional Subdivisions and Processing Streams.

Hermes D, Siero JC, Aarnoutse EJ, Leijten FS, Petridou N, Ramsey NF (2012) Dissociation between neuronal activity in sensorimotor cortex and hand movement revealed as a function of movement rate. The Journal of neuroscience : the official journal of the Society for Neuroscience 32:9736–9744.

Hikosaka O, Nakamura K, Sakai K, Nakahara H (2002) Central mechanisms of motor skill learning. Current opinion in neurobiology 12:217–222.

Honda M, Deiber MP, Ibáñez V, Pascual-Leone A, Zhuang P, Hallett M (1998) Dynamic cortical involvement in implicit and explicit motor sequence learning. A PET study. Brain 121:2159–2173.

Hutton C, Bork A, Josephs O, Deichmann R, Ashburner J, Turner R (2002) Image distortion correction in fMRI: a quantitative evaluation. Neuroimage 16:217240.

Indovina I, Sanes JN (2001) On somatotopic representation centers for finger movements in human primary motor cortex and supplementary motor area. Neuroimage 13:1027–1034.

Karni A, Meyer G, Jezzard P, Adams MM, Turner R, Ungerleider LG (1995) Functional MRI evidence for adult motor cortex plasticity during motor skill learning. Nature 377:155–158.

Karni A, Meyer G, Rey-Hipolito C, Jezzard P, Adams MM, Turner R, Ungerleider LG (1998) The acquisition of skilled motor performance: fast and slow experience-driven changes in primary motor cortex. Proc Natl Acad Sci U S A 95:861–868.

Kass RE, Raftery AE (1995) Bayes Factors. Journal of the American Statistical Association 90:773–795.

Kawashima R, Matsumura M, Sadato N, Naito E, Waki A, Nakamura S, Matsunami K, Fukuda H, Yonekura Y (1998) Regional cerebral blood flow changes in human brain related to ipsilateral and contralateral complex hand movements-a PET study. European Journal of Neuroscience 10:2254–2260.

Kornysheva K, Diedrichsen J (2014) Human premotor areas parse sequences into their spatial and temporal features. Elife 3:e03043.

Laje R, Buonomano DV (2013) Robust timing and motor patterns by taming chaos in recurrent neural networks. Nature neuroscience 16:925–933.

Lashley KS (1951) The problem of serial order in behavior. In: Cerebral mechanisms in behavior, pp 112–136.

Lawrence DG, Kuypers HG (1968) The functional organization of the motor system in the monkey. I. The effects of bilateral pyramidal lesions. Brain 91:1–14.

Logothetis NK, Pauls J, Augath M, Trinath T, Oeltermann A (2001) Neurophysiological investigation of the basis of the fMRI signal. Nature 412:150–157.

Matsuzaka Y, Picard N, Strick PL (2007) Skill representation in the primary motor cortex after long-term practice. J Neurophysiol 97:1819–1832.

Muir RB, Lemon RN (1983) Corticospinal neurons with a special role in precision grip. Brain research 261:312–316.

Mushiake H, Inase M, Tanji J (1991) Neuronal activity in the primate premotor, supplementary, and precentral motor cortex during visually guided and internally determined sequential movements. Journal of Neurophysiology 66:705–718.

Oosterhof NN, Wiestler T, Downing PE, Diedrichsen J (2011) A comparison of volume-based and surface-based multi-voxel pattern analysis. Neuroimage 56:593–600.

Picard N, Matsuzaka Y, Strick PL (2013) Extended practice of a motor skill is associated with reduced metabolic activity in M1. Nature neuroscience 16:1340–1347.

Sakai K, Hikosaka O, Miyauchi S, Takino R, Sasaki Y, Pütz B (1998) Transition of brain activation from frontal to parietal areas in visuomotor sequence learning. Journal of Neuroscience 18:1827–1840.

Schieber MH, Hibbard LS (1993) How somatotopic is the motor cortex hand area? Science 261:489–492.

Shima K, Tanji J (1998) Both supplementary and presupplementary motor areas are crucial for the temporal organization of multiple movements. J Neurophysiol 80:3247–3260.

Siero JC, Hermes D, Hoogduin H, Luijten PR, Petridou N, Ramsey NF (2013) BOLD consistently matches electrophysiology in human sensorimotor cortex at increasing movement rates: a combined 7T fMRI and ECoG study on neurovascular coupling. J Cereb Blood Flow Metab 33:1448–1456.

Stephan KE, Penny WD, Daunizeau J, Moran RJ, Friston KJ (2009) Bayesian model selection for group studies. Neuroimage 46:1004–1017.

Tanji J, Shima K (1994) Role for supplementary motor area cells in planning several movements ahead. Nature 371:413–416.

Walther A, Nili H, Ejaz N, Alink A, Kriegeskorte N, Diedrichsen J (2016) Reliability of dissimilarity measures for multi-voxel pattern analysis. Neuroimage 137:188–200.

Wiestler T, Diedrichsen J (2013) Skill learning strengthens cortical representations of motor sequences. Elife 2:e00801.

Wiestler T, McGonigle DJ, Diedrichsen J (2011) Integration of sensory and motor representations of single fingers in the human cerebellum. J Neurophysiol 105:3042–3053.

Wiestler T, Waters-Metenier S, Diedrichsen J (2014) Effector-independent motor sequence representations exist in extrinsic and intrinsic reference frames. The Journal of neuroscience : the official journal of the Society for Neuroscience 34:5054–5064.

